# Brain I3 Binding Protein regulates K-Ras4B membrane localization and signaling

**DOI:** 10.1101/2021.09.21.461216

**Authors:** Ian McCabe, Huanqing Zhang, Jonathan A. Cooper, David L. Turner, Anne B. Vojtek

**Affiliations:** Department of Biological Chemistry, University of Michigan, Ann Arbor, MI 48109-0600; Molecular Neuroscience Institute, University of Michigan, Ann Arbor, MI 48109; Fred Hutchinson Cancer Research Center, Seattle, WA 98109

## Abstract

Membrane localization of Ras proteins is necessary for their biological functions and oncogenic activity. We report here on the identification of Brain I3 Binding Protein (BRI3BP) as a novel binding partner for Ras. We show that K-Ras4B plasma membrane localization and biological function are reduced in the absence of BRI3BP. BRI3BP interacts with K-Ras4B and K-Ras4A and our data suggest that BRI3BP operates within the recycling endosomal compartment to regulate K-Ras localization to the plasma membrane. This study uncovers a new regulatory protein for Ras membrane localization.

## Introduction

Ras proteins are central regulators of cell proliferation that operate as molecular switches, cycling between GTP-active and GDP-inactive states (1, 2). This transition between active and inactive states is tightly regulated by guanine nucleotide exchange factors (GEFs) and GTPase activating proteins (GAPs). GEFs promote the formation of GTP, active Ras whereas GAPs accelerate the intrinsic GTP hydrolytic activity of Ras to inactivate Ras. GTP-Ras regulates cell proliferation by activation of downstream effector pathways, most prominent among which is the Raf-Mek-Erk mitogen activated protein kinase (MAPK) cascade (3, 4).

Four mammalian Ras proteins, K-Ras4A, K-Ras4B, H-Ras and N-Ras, are encoded by three RAS genes, with K-Ras4A/4B generated by alternate splicing from the K-RAS locus (5). Ras mutant proteins, with mutations at G12, G13 and Q61, exhibit impaired intrinsic and GAP stimulated GTP hydrolytic activity, resulting in prolonged, activated Ras signaling and activation of downstream effector pathways (1, 2). K-Ras mutations are prevalent in three of the four most lethal human cancers, pancreatic, lung and colon cancers (1, 6). These activating mutations in Ras drive tumor initiation and progression, and resistance to cancer therapy is frequently associated with increased Ras activity, either by mutation or amplification (1, 6, 7).

Ras proteins localize to the inner surface of the plasma membrane where they recruit downstream effectors and activate signaling pathways (8, 9). A C-terminal membrane anchor targets Ras to the plasma membrane. The membrane anchor is comprised of at least two signals that act in concert. The first signal, common to all Ras isoforms, is a farnesyl carboxyl methyl ester (10). This post-translational modification of the C-terminal CAAX motif of Ras is the product of three sequential reactions, which are catalyzed by cytosolic farnesyl transferase, endoplasmic reticulum (ER)-localized Ras converting enzyme, and isoprenyl cysteine carboxyl methyl transferase. A second signal for membrane attachment, common to H-Ras and N-Ras, is palmitoylation, which takes place at one or two cysteines adjacent to the farnesyl methyl ester (10). For K-Ras4B, the second component of the membrane anchor is not a hydrophobic lipid but instead is a polybasic domain located adjacent to the farnesyl methyl ester (11, 12). The polybasic domain binds with high affinity to phosphatidylserine, which is enriched at the cytosolic surface of the plasma membrane and recycling endosomes (13–16). In addition to the farnesyl carboxyl methyl ester, K-Ras4A, which has a split polybasic domain, is palmitoylated (17). These multi-component Ras membrane anchors are required and sufficient for plasma membrane association. Ras membrane association is central to transformation and oncogenicity, and Ras proteins that fail to associate with the plasma membrane are not oncogenic (8, 9).

The spatial distribution of Ras in cellular membranes and localization to the plasma membrane is a regulated process, with phosphodiesterase 6 delta (PDEδ) and G-protein coupled receptor 31 (GPR31) playing central roles (18–21). PDEδ facilitates the trafficking of Ras proteins from endomembranes to the recycling endosomal compartment, whereas GPR31 facilitates the trafficking of Ras proteins from the ER to the plasma membrane. The interaction of Ras with PDEδ and GPR31 is dependent on farnesylation (prenylation) of the Ras membrane anchor. Additionally, calmodulin interacts with farnesylated/prenylated K-Ras. The calmodulin interaction has different roles depending on cellular context, extracting K-Ras from the plasma membrane or activating phosphatidylinositol 3-kinase/Akt in the presence of the signaling lipids phosphatidylinositol 4, 5 bisphosphate and phosphatidylinositol 5-phosphate (22, 23).

In this current study, we report the identification of BRI3BP as an interaction partner for Ras. In mammalian cells, BRI3BP interacts with K-Ras, but not H-Ras, and regulates K-Ras membrane localization and transforming activity. Like PDEδ and GPR31, the interaction of BRI3BP with Ras is dependent on farnesylation of the membrane anchor. The localization of BRI3BP to the recycling endosomal compartment suggests that BRI3BP operates within this cellular compartment to regulate K-Ras localization to the plasma membrane.

## Results

Multiple Ras interacting proteins (Rips) were identified in a yeast two-hybrid screen, including c-Raf (Rip51), phosphatidylinositol 3-kinase δ, and Ral guanine nucleotide dissociation stimulator (24–27). c-Raf is a Ras effector protein and mediates Ras signaling, and mutations in the Ras effector domain abolish the interaction of Ras with c-Raf/Rip51 (26–28). Rip80 was identified as a Ras interacting protein in the same yeast two hybrid screen but not studied further until this report. In contrast to c-Raf, Rip80 interacts with Ras in an effector domain independent manner (Figure S1A). In addition, the interaction of Rip80 with Ras is dependent on prenylation, as mutation of the cysteine within the Ras CAAX box abolishes binding to Ras (Figure S1B). Rip80 encodes amino acids 85–253 of BRI3BP, which was initially discovered as a binding protein for brain I3 (BRI3), a poorly characterized TNFa upregulated protein in brain endothelial cells (29). Ras function is regulated by GAPs, GEFs, and prenyl recognition proteins. The latter class of regulatory proteins bind to post-translationally modified prenylated Ras proteins and regulate Ras membrane association. The yeast two-hybrid interaction profile of Rip80 with Ras suggests that BRI3BP may function as a Ras prenyl recognition protein.

To test for co-association of BRI3BP and Ras in mammalian cells, pull-down assays were performed. For these studies, we generated a human HEK293 cell line in which an HA epitope tag was added to the C-terminus of the endogenous BRI3BP protein using CRISPR/Cas9 and homology directed repair (HDR). This cell line was transfected with expression vectors for GST (control) or GST tagged constitutively activated K-Ras4B (GST-K-RasV12) and pull-down assays performed. Endogenous BRI3BP-HA co-associates with activated GST-K-Ras4B but not GST control (Figure 1A). In addition, consistent with the yeast two-hybrid results, the interaction between BRI3BP and K-Ras4B in mammalian cells is dependent on Ras prenylation, as mutation of the <u>C</u>AAX box to <u>S</u>AAX abolishes the interaction (Figure 1A). To determine if BRI3BP demonstrates a preference for GDP-vs GTP-bound Ras, we compared the binding of wild type K-Ras4B after serum starvation (GDP-Ras) or starvation plus serum stimulation for 10 minutes (GTP-Ras). K-Ras4B bound equally well to BRI3BP under either condition, indicating that the interaction between K-Ras4B and BRI3BP is nucleotide independent (Figure 1B).

**Figure 1.**
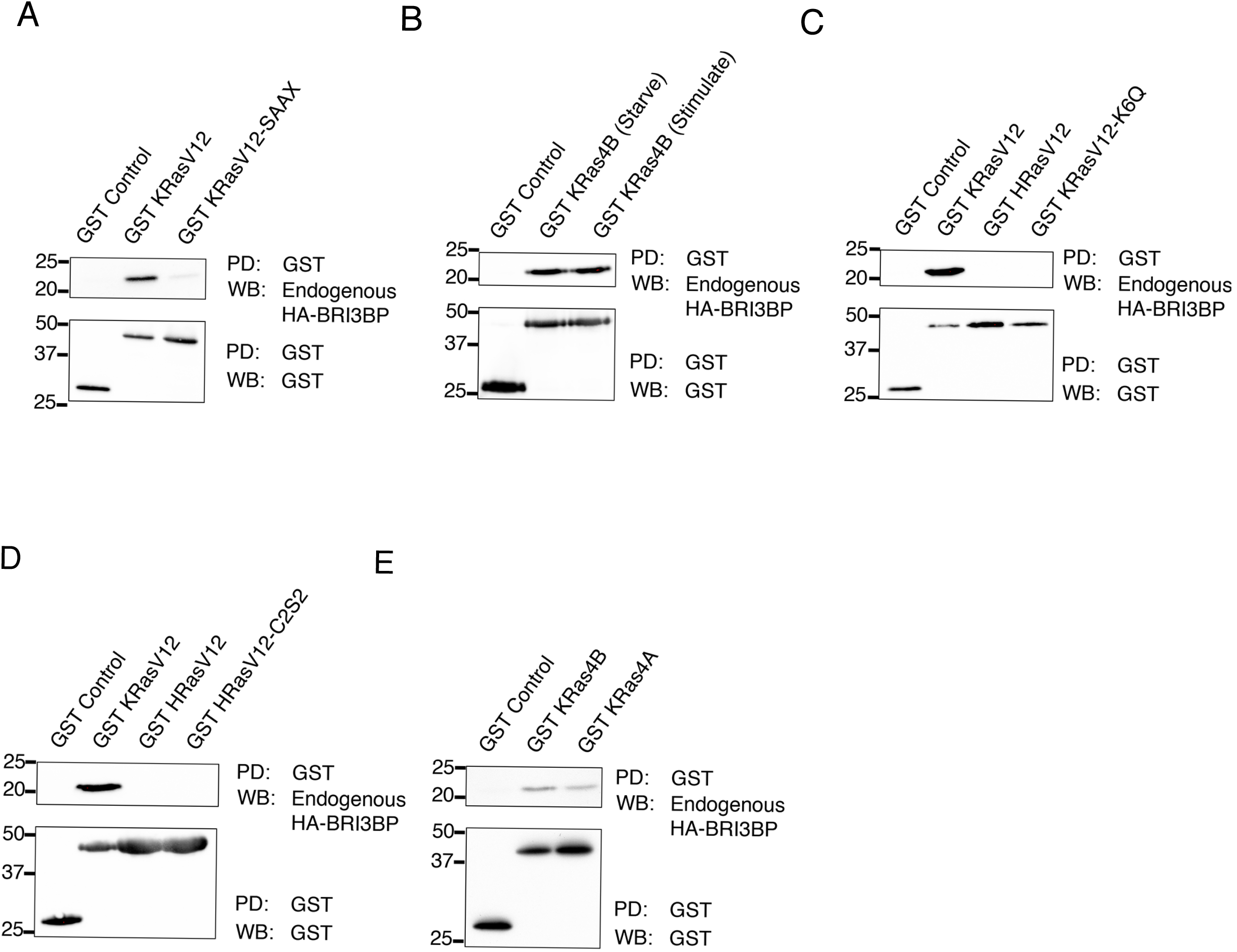
BRI3BP interacts with K-Ras4B and K-Ras4A. GST control or GST-fusion proteins were expressed in HEK cells in which the endogenous BRI3BP protein has been engineered to include a C-terminal HA epitope tag. GST or GST fusion proteins were isolated (or pulled-down, PD) from extracts with glutathione magnetic beads and western blotted (WB) for co-associating BRI3BP-HA. A. BRI3BP interacts with K-Ras4B-V12 and the interaction depends on prenylation as the K-Ras4B-V12-SAAX mutant does not interact with BRI3BP. B. BRI3BP co-associates with wild-type K-Ras4B and the interaction is not modulated by serum starvation, indicating the interaction is independent of nucleotide bound to K-Ras4B. C. BRI3BP interacts with K-Ras4B-V12 but not H-Ras-V12, and the polylysine motif in the HVR of K-Ras4B is required for association. D. H-Ras-V12-C2S2 also fails to interact with BRI3BP, indicating that palmitoylation of H-Ras does not block binding to BRI3BP. E. The K-Ras4A isoform also co-associates with BRI3BP.

H-Ras and K-Ras4B exhibit 85% overall sequence identity and thus share effectors and regulators of the GTP/GDP cycle, including GAPs and GEFs (1, 6). The two proteins diverge significantly in the C-terminal hypervariable region (HVR) that regulates membrane localization, although both H- and K-Ras have in common a C-terminal farnesyl methyl ester (1, 9, 11). To determine if BRI3BP interacts with H-Ras in addition to K-Ras4B in mammalian cells, constitutively active H-Ras (GST-H-RasV12) was expressed in BRI3BP-HA tagged HEK cells and co-association assessed. In striking contrast to K-Ras4B, H-Ras does not interact with endogenous BRI3BP-HA (Figure 1C). The selective binding of BRI3BP to K-Ras4B but not H-Ras in mammalian cells was initially surprising because H-Ras was the bait in the yeast two-hybrid screen. We tested whether co-association could be detected with the BRI3BP domain identified in the two-hybrid assay. Even under conditions in which the BRI3BP domain is overexpressed, a striking selectivity (~7-fold) for the K-Ras isoform is observed (Figure S2). We conclude that BRI3BP is a preferential binding partner for K-Ras4B when BRI3BP is expressed at normal, endogenous levels.

The HVR plays a critical role in Ras membrane localization and is the most divergent region between K- and H-Ras (1, 8, 9). A distinguishing feature of the K-Ras4B HVR is the polybasic region. To gain insight into the molecular basis for the discrimination in binding of the Ras proteins by BRI3BP in mammalian cells, the polybasic region of K-Ras was mutated, with 6 lysines changed to electrostatically neutral glutamine residues (K6/Q). Mutation of the polybasic residues within the HVR of K-Ras abolished binding to BRI3BP (Figure 1C), consistent with the selective binding of BRI3BP to K-Ras but not H-Ras. In addition to lacking the basic residues, H-Ras also differs from K-Ras4B in palmitoylation at two cysteines in the HVR (10, 11). Palmitoylation is a dynamic post-translational lipid modification that together with farnesylation regulates H-Ras membrane localization (30, 31). To determine if palmitoylation of H-Ras interferes with recognition by BRI3BP, a palmitoylation defective H-Ras was generated in which the two sites of palmitoylation were mutated and its interaction with BRI3BP tested by pull-down analysis. Neither H-Ras nor the palmitoylation deficient H-Ras (H-RasV12 C2S2) interact with BRI3BP, indicating that palmitoylation of H-Ras does not prevent its binding to BRI3BP (Figure 1D).

The K-Ras locus encodes two different Ras proteins, K-Ras4B and K-Ras4A, due to alternate splicing of exon 4. Exon4A and 4B encode the HVR region of K-Ras4A and K-Ras4B, respectively. Because activating mutations in Ras most often occur in exon 1 or 2, both K-Ras4A and 4B are oncogenic (17, 32–34). To assess BRI3BP binding, GST-K-Ras4A was expressed in BRI3BP-HA cells and co-association was determined. BRI3BP interacts with K-Ras4A (Figure 1E).

Oncogene transfected NIH3T3 mouse fibroblasts undergo morphological transformation and loss of contact inhibition, leading to the formation of multilayered foci (35). To determine the role of BRI3BP in regulating the transforming potential of oncogenic K-Ras, focus formation assays in NIH3T3 cells were performed. Control or BRI3BP RNAi expression vectors were transfected into NIH3T3 cells together with an expression vector for activated K-Ras4B, and focus formation assessed 21 days later. Oncogenic K-RasV12 efficiently induced focus formation in NIH3T3 cells. Significantly, oncogenic K-RasV12-driven focus formation was decreased after RNAi mediated reduction of BRI3BP function (Figure 2), indicating that BRI3BP function is required for K-RasV12 transforming activity in NIH3T3 cells.

**Figure 2.**
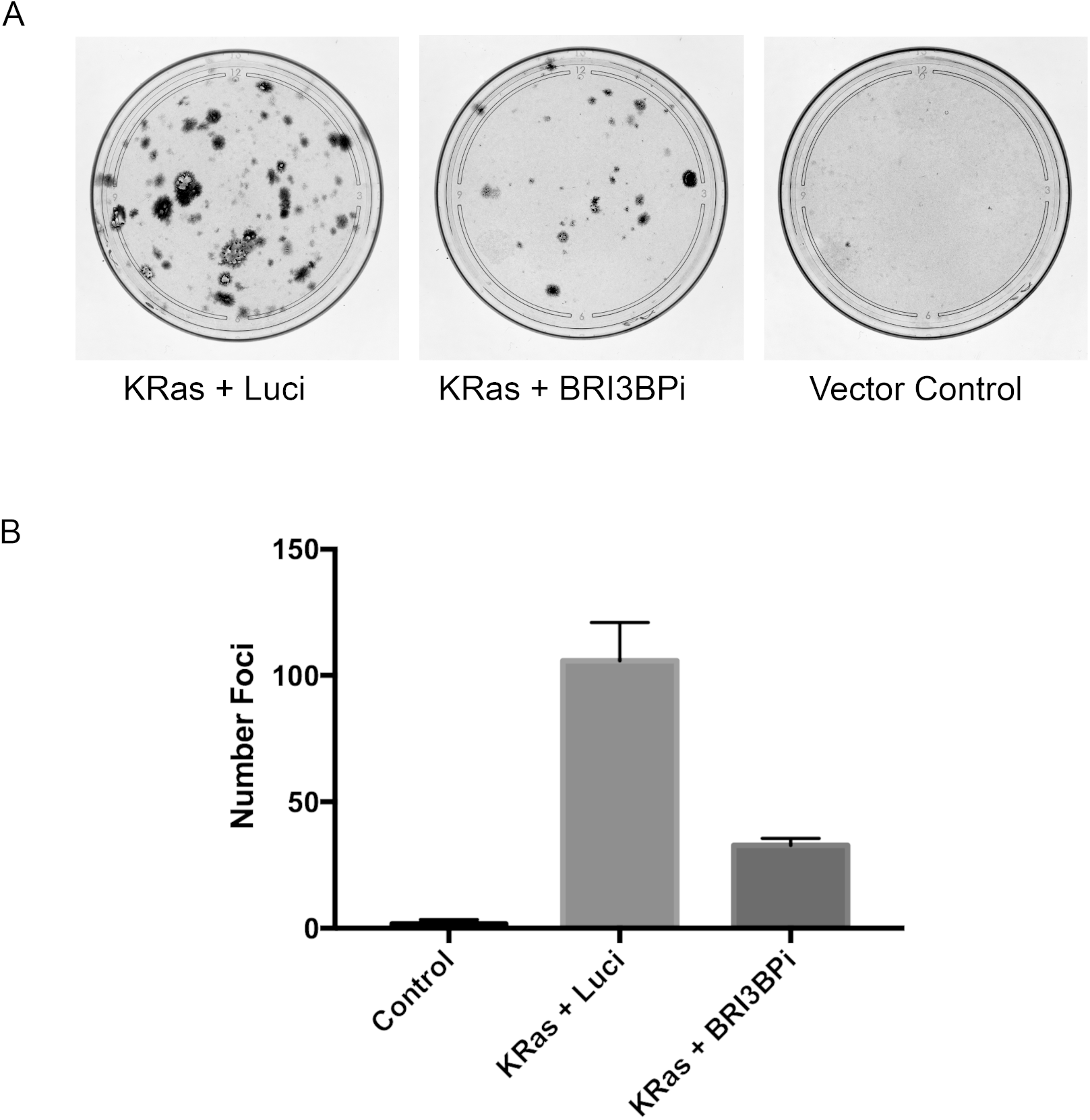
BRI3BP regulates oncogenic K-Ras transforming activity. NIH3T3 cells were cotransfected with a K-Ras4B-V12 expression vector or a control expression vector and a luciferase RNAi (Luci) control vector or a BRI3BP RNAi (BRI3BPi) vector. A. Foci visualized by crystal violet staining after fixation at 21 days. RNAi against BRI3BP reduced foci formation by K-Ras4B-V12 relative to luciferase control. B. Quantitation of foci (average ± standard deviation).

Given the dependency of the BRI3BP-K-Ras4B interaction on the K-Ras4B membrane anchor (Figure 1A, C), we investigated whether BRI3BP regulates membrane localization of K-Ras4B. To assess the role of BRI3BP, we generated HeLa cells in which the endogenous BRI3BP genes were disrupted using CRISPR/Cas9. Because of the lack of antibodies that specifically and selectively recognize K-Ras (but not H- or N-Ras), we introduced a vector that expresses a green fluorescent protein (GFP) K-Ras4B fusion protein (GFP-K-Ras4B-V12) into wild type or BRI3BP knockout HeLa cells to examine K-Ras4B localization. GFP-K-Ras4B localization was analyzed in fixed, stained cells by confocal imaging. Knockout of BRI3BP causes mislocalization of K-Ras4B, with knockout cells showing decreased localization of K-Ras4B at the plasma membrane, relative to wild type cells (Figure 3A, localization of GFP-K-Ras in wild type (WT) cells, upper left panel, compared to BRI3BP knockout (KO) cells, lower left panel). Representative images are shown in Figure 3A, with additional images in Figure S3.

**Figure 3.**
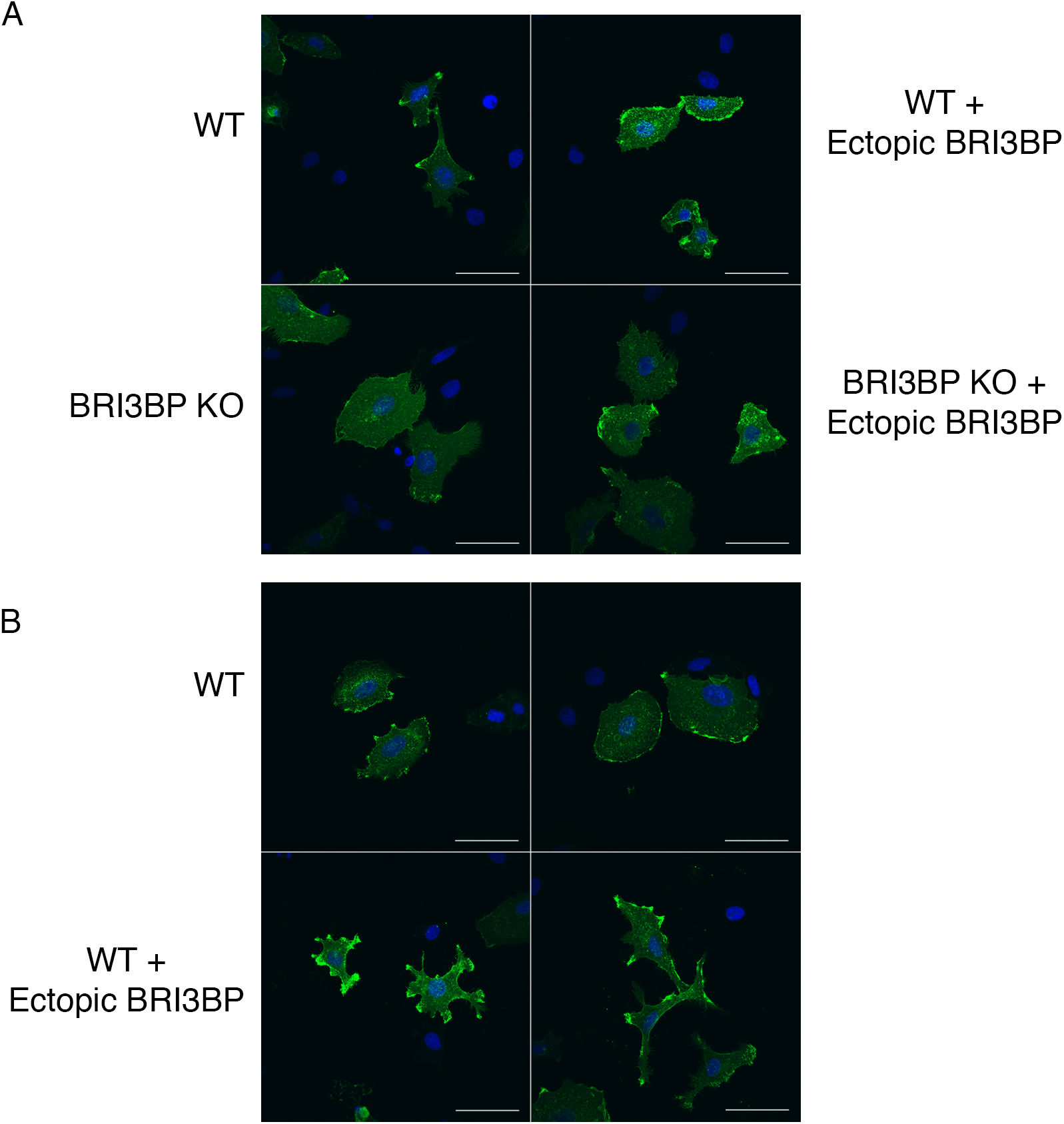
Loss of BRI3BP reduces K-Ras4B plasma membrane localization. A. Confocal images of GFP-K-Ras4B-V12 localization in wild-type HeLa cells (WT, upper left panel), WT cells with overexpressed BRI3BP from a transfected expression vector (WT + ectopic BRI3BP, upper right panel), CRISPR/Cas9 BRI3BP knockout HeLa cells (KO, lower left panel), or KO cells with ectopic BRI3BP from a transfected expression vector to allow complementation (KO + ectopic BRI3BP, lower right panel). B. GFP-K-Ras4B-V12 localization in wild type cells with (lower two panels) and without (upper two panels) ectopic BRI3BP. Cells were fixed at 48 hours (A) or 72 hours (B) after transfection, then processed for indirect immunofluorescent detection of GFP. Scale bar: 50 μm.

Importantly, normal localization of K-Ras4B in the BRI3BP knockout line is restored by expression of BRI3BP from a cDNA expression vector, confirming that K-Ras mislocalization is due to loss of BRI3BP function in these cells (Figure 3A, lower right panel). The decrease in plasma membrane localization is not an indirect effect of decreased stability or increased degradation of K-Ras, as levels of GFP-K-Ras are similar in both wild type and knockout cells, based on western blot analysis (Figure S4). Finally, overexpression of BRI3BP in wild type HeLa cells enhances plasma membrane localization of GFP-K-Ras4B (Figure 3A, upper right panel, and Figure 3B). Taken together, the loss of function and overexpression data indicate that BRI3BP regulates K-Ras4B localization to the plasma membrane.

To further investigate the relationship between K-Ras and BRI3BP, we determined the subcellular localization of BRI3BP and investigated whether BRI3BP co-localizes with K-Ras. Because of the paucity of BRI3BP antibodies and, as noted above, antibodies that specifically and selectively recognize K-Ras but not H- or N-Ras, we transiently co-transfected HeLa cells with expression vectors for GFP-K-Ras4B-V12 and BRI3BP fused to mScarlet (36). BRI3BP localized predominantly to a perinuclear region of the cell, and GFP-K-Ras4B and BRI3BP-mScarlet co-localized within this compartment (Figure 4A). BRI3BP-mScarlet did not localize to the plasma membrane and no co-localization of GFP-K-Ras4B-V12 and BRI3BP-mScarlet was apparent at the plasma membrane.

**Figure 4.**
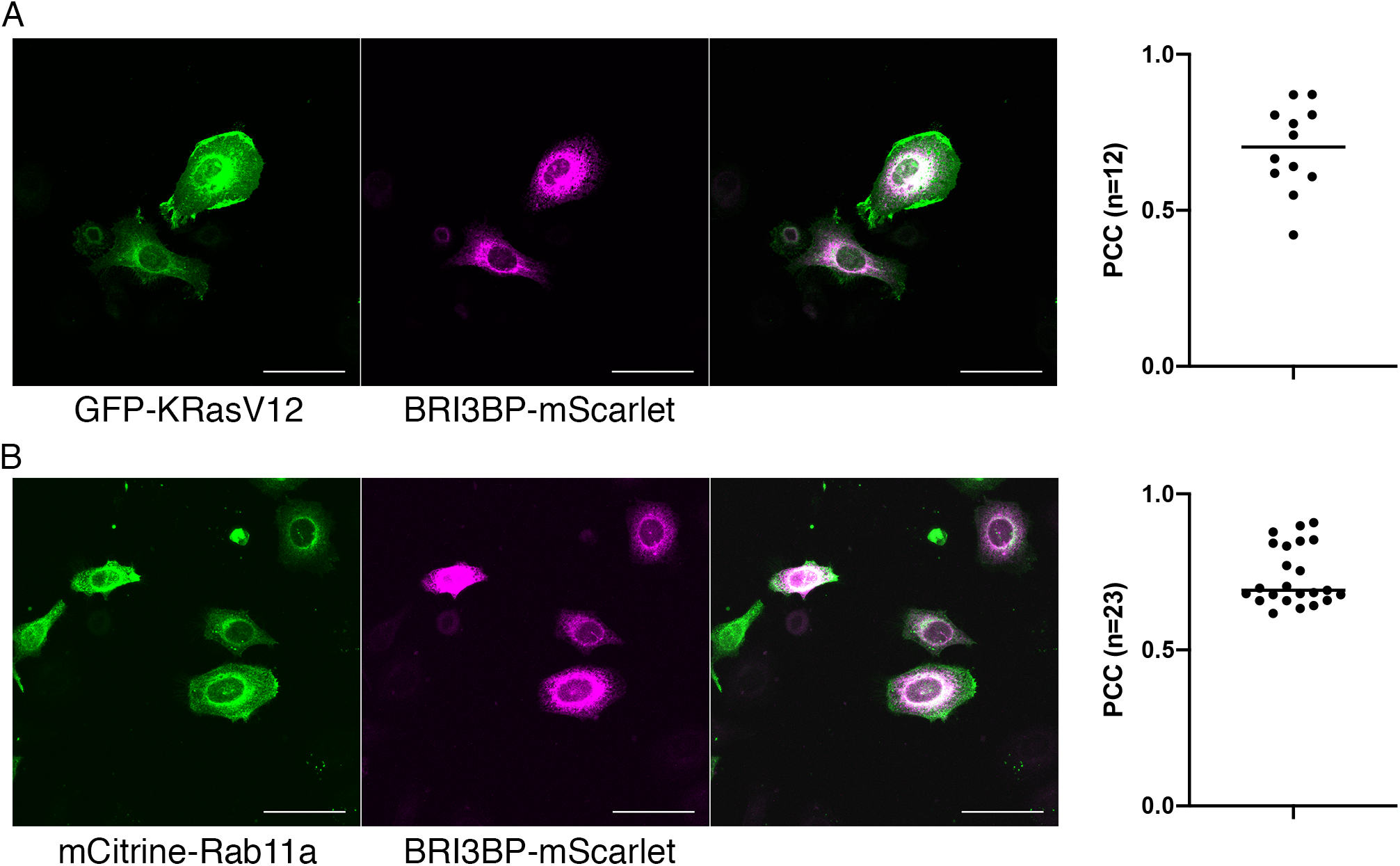
BRI3BP co-localizes with K-Ras4B and the recycling endosomal marker Rab11a. Confocal images of fixed HeLa cells transfected with GFP-K-RasV12 and BRI3BP-mScarlet (A) or BRI3BP-mScarlet and mCitrine-Rab11a, a marker of the recycling endosomal compartment (B). Single labeling is shown in left and middle panels, merged images in right panel. Quantitation of colocalization, with Pearson’s correlation coefficients between GFP-KRasV12 and BRI3BP-mScarlet in (A) and mCitrine-Rab11a and BRI3BP-mScarlet in (B). Each dot represents a single cell.

K-Ras4B traffics to the plasma membrane via the perinuclear recycling endosomal compartment (12, 17, 37). To investigate potential molecular mechanisms for BRI3BP in regulation of K-Ras plasma membrane localization, and given the observed perinuclear localization of BRI3BP, we asked whether BRI3BP overlaps with Rab11a, a marker of the recycling endosomal compartment (38–40). Previous reports have shown that K-Ras and Rab11a overlap in the perinuclear recycling endosome (12). Using confocal microscopy of fixed cells, we observe substantial overlap between BRI3BP-mScarlet and mCitrine-Rab11a (Figure 4B). We next compared localization of BRI3BP against Rab5a, a marker of early endosomes. No significant co-localization was observed between BRI3BP and Rab5a (Figure S5). Taken together, these observations are consistent with BRI3BP functioning within the recycling endosomal compartment to enable K-Ras enrichment at the plasma membrane.

## Discussion

We identify BRI3BP as a new binding partner for Ras and demonstrate that BRI3BP is required for K-Ras4B localization and signaling. We find that the interaction of K-Ras4B with BRI3BP depends on K-Ras prenylation and is independent of the nucleotide bound to Ras, consistent with BRI3BP acting as a prenyl recognition protein for K-Ras. In mammalian cells, endogenous BRI3BP interacts with K-Ras4B but not H-Ras. The two Ras proteins have distinct hypervariable regions that regulate membrane localization (1, 8, 9). Consistent with the selectivity of BRI3BP for K-Ras4B, we demonstrate a requirement for the K-Ras4B polybasic domain, which is not present in H-Ras.

Two recent reports provide independent support for a role for BRI3BP in K-Ras signaling. BRI3BP was identified in a recent mass spectrometry analysis of endogenous proteins that co-immunoprecipitated with K-Ras [Supplemental TableS2 in (41)]. BRI3BP coimmunoprecipitated with K-Ras in pancreatic and lung cancer cell lines that express different KRas alleles frequently mutated in human cancers, G12V, G12D and G12C (41)]. Although the BRI3BP/K-Ras interaction was not studied further in that report, the mass spectrometry data provides independent confirmation of the observation reported here that BRI3BP is a Ras binding protein. In addition, different K-Ras alleles (G12V, G12D, G12R) have unique signaling properties (42–44). K-Ras G12R cells, for example, exhibit impaired/reduced phosphatidylinositol 3-kinase signaling (45). Intriguingly, BRI3BP was identified as a K-Ras G12R allele-specific genetic dependency in CRISPR/Cas9 knockout analyses of cancer cell lines, providing further support for a role for BRI3BP in K-Ras signaling (46).

Ras signals at the plasma membrane, and we show here that BRI3BP is required for K-Ras4B localization. In BRI3BP knockout cells, plasma membrane localization of K-Ras4B is reduced. This phenotype is complemented by expression of BRI3BP, demonstrating that mislocalization of K-Ras in the knockout cells is due to loss of BRI3BP function. Additionally, in wild type cells that ectopically express BRI3BP, K-Ras localization at the plasma membrane is increased. The complementary phenotypes observed with loss of function and overexpression (decreased or increased Ras plasma membrane localization, respectively) provide compelling support for BRI3BP as a regulatory protein for K-Ras4B localization. Membrane association is a key event in Ras transformation and oncogenicity, and Ras proteins that fail to localize to the plasma membrane are neither active nor oncogenic (8, 9). Concordant with K-Ras4B mislocalization in BRI3BP knockout cells, focus formation, a cellular measure of oncogenic KRas activity, is decreased when BRI3BP function is reduced in NIH3T3 cells.

K-Ras is transported to the plasma membrane from the perinuclear recycling endosome (12). Co-localization of BRI3BP with Rab11a, a marker of the perinuclear recycling endosomal compartment (38–40), but not the early endosomal marker Rab5a, suggests that BRI3BP operates within the recycling endosomal compartment to regulate K-Ras plasma membrane localization. The cytosolic surface of the recycling endosome is negatively charged, similar to the inner surface of the plasma membrane. Electrostatic interactions and high affinity binding to phosphatidylserine by the K-Ras4B polybasic domain have been proposed as a mechanism to trap K-Ras in the recycling endosome, prior to vesicular transport to the plasma membrane (12, 16). We propose that the interaction of the K-Ras4B prenylated hypervariable region with BRI3BP and the polybasic domain with negatively charged membrane phospholipids within the recycling endosomal compartment operate together to facilitate the trafficking of the K-Ras4B protein to the plasma membrane. Mutational analysis of the K-Ras4B hypervariable region provides support for this model.

The Ras prenyl recognition protein Pdeδ is a cytosolic chaperone that assists in the intracellular transport of Ras proteins from endomembranes to the perinuclear recycling endosomal compartment. At the recycling endosomal compartment, the small GTPase Arl-2 facilitates the off-loading of Ras from Pdeδ (12, 18). BRI3BP, present in the recycling endosomal compartment, is positioned spatially to operate downstream of Pdeδ, and future studies will be directed at determining the relationship between BRI3BP and Pdeδ in regulating K-Ras trafficking. The G-protein coupled receptor GPR31 also regulates membrane association of K-Ras, acting as a secretory pathway chaperone for trafficking Ras from the endoplasmic reticulum to the plasma membrane (20). The role of GPR31 and BRI3BP, whether separate or cooperative for Ras localization, is not known. It seems likely that K-Ras spatial organization is regulated by a network of regulatory proteins, including BRI3BP. Combinatorial loss of function studies to examine the interplay among BRI3BP and prenyl recognition protein pathways are likely to provide important insights into molecular mechanisms for Ras trafficking.

BRI3BP interacts with both K-Ras4B and K-Ras4A. In contrast to H-Ras, neither K-Ras4B nor K-Ras4A traffic through the Golgi to reach the plasma membrane (17, 37). Thus, BRI3BP, operating from a site of action within the recycling endosomal compartment, may select K-Ras4B and K-Ras4A but not H-Ras for trafficking to the plasma membrane, thereby preventing their accumulation and/or trafficking to the Golgi compartment, the fate of H-Ras.

Membrane localization of Ras proteins are necessary for both normal cellular functions and for oncogenesis. Our studies identify BRI3BP as a regulatory step for K-Ras membrane localization and function. The selectivity of BRI3BP for K-Ras but not H-Ras provides an opportunity for future targeted therapies that block oncogenic K-Ras without targeting H-Ras, potentially decreasing toxicity from blocking Ras function in its entirety.

## Methods

### Cell Lines

Human embryonic kidney (HEK) 293 and HeLa cells were from ATCC and were grown in Dulbecco’s Modified Eagle Medium (DMEM) with high glucose and sodium pyruvate (Gibco#11995065) supplemented with 10% fetal bovine serum (Gibco#26140079) and penicillin-streptomycin (Gibco#15140122). NIH3T3 cells, from Channing Der, were grown in DMEM supplemented with 10% calf serum (Hyclone). Cell lines were grown at 37°C with 5% CO_2_.

HEK cells were transfected with calcium phosphate (Promega, Profection Mammalian Transfection System), HeLa cells with HeLa monster (Mirus), and NIH3T3 cells with Trans-IT X2 (Mirus), according to the manufacturer’s instructions.

HEK 293 BRI3BP-HA contains an HA epitope tag inserted into the C-terminus of BRI3BP. The cell line was generated by CRISPR/Cas9 and homology directed repair (HDR). Primers for cloning into pX458 (47): BRI3BP gRNA 3F, 5’-CACCGGCTGACCTTCACTTGTCCT-3’ and BRI3BP gRNA 4R, 5’-AAACAGGACAAGTGAAGGTCAGCC-3’. The pX458 3/4 sgRNA cuts 5 nucleotides upstream of the BRI3BP stop codon. The BRI3BP GSG HA Tag insertion donor used for HDR was a single stranded DNA oligo donor (171 nts, IDT) with two phosphorothioate linkages at each end, a Gly-Ser-Gly linker upstream of the HA epitope, and asymmetric homology arms to BRI3BP. To confirm fusion of the HA-tag to BRI3BP, the HEK 293 BRI3BP-HA cell line was transfected with control or BRI3BP silencer select siRNAs with Lipofectamine RNAi Max (ThermoFisher Scientific), according to manufacturer’s instructions. Silencer select siRNAs (ThermoFisher Scientific): Negative control siRNA, Catalog # 4390843; BRI3BP siRNA oligo1 (Catalog # 4392420, siRNA ID#s44364); and, BRI3BP siRNA oligo 2 (4392420, siRNA ID# s44362). Western blot analysis with an anti-HA antibody (Cell Signaling Technology, monoclonal rabbit antibody #3724) of extracts prepared 48 or 72 hours after transfection was used to detect the level of the BRI3BP-HA fusion protein. Endogenous BRI3BP-HA levels were reduced in cells transfected with two BRI3BP siRNAs targeting different sequences relative to the control siRNA, confirming fusion of the HA epitope tag to BRI3BP (Fig. S6, n=2).

The HeLa BRI3BP knockout cell line was constructed using CRISPR/Cas9. Two pX458 vectors with sgRNAs targeting exon2 or exon3 were co-transfected together with a vector that expresses a puromycin resistance gene, included at 10% of the total plasmid DNA transfected. Transfected cells were selected with puromycin (2 μg/ml). The presence of the deletion was confirmed in the puromycin selected population then single cells clones were isolated by limiting dilution. Genomic DNA was prepared from single cell clones and deletions/indels identified by PCR and sequence analysis of genomic DNA. As expected, sequence analysis of HeLa #18 with primers spanning the two gRNA cut sites identifies a deletion between the sgRNA target sites in exon 2 and 3 in the BRI3BP gene. The line also has an allele with an indel in exon2 (a 2 nucleotide deletion, CRISPR-ID analysis). No wild type exon 2 alleles remain.

Primers for cloning into pX458 to generate the exon 2 targeting sgRNA: BRI3BP7a, 5’-CACCGATATGTTCGTGGAGACACTG-3’ and BRI3BP7b, 5’-AAACCAGTGTCTCCACGAACATATC-3’. pX458 sgRNA 3/4 which targets exon 3 is described above.

### Yeast Two-hybrid Assay

The Saccharomyces cerevisiae L40 reporter strain was transformed with expression vectors for LexA DNA binding domain and VP16 activation domain fusion proteins (26). Expression vectors for LexA DNA binding domain Ras and Ras mutant fusion proteins have been described (48). VP16-Rip80 was identified as a Ras interacting partner in a two-hybrid screen of a mouse embryo cDNA library (26) and encodes amino acids 85 to 253 of BRI3BP. Yeast were transformed and growth assays in minimal media lacking histidine were performed to assess activation of the His3 reporter. Interaction of two hybrid fusion proteins in this system leads to activation of the His3 reporter and supports growth in minimal media in the absence of histidine.

### GST Pull Down Assays

HEK 293 BRI3BP-HA cells were transfected with a pEBG vector expressing GST control or GST-Ras fusion proteins, wild type or mutant. In pEBG, a GST fusion protein is expressed in mammalian cells under control of an EF1alpha promoter (49).

GST is fused to the N-terminus of the Ras protein. Per pull-down, 3 wells of a 12-well dish were each transfected with 1 μg DNA. 24 hours after transfection, extracts were prepared in Tris Lysis Buffer (25 mM Tris-HCl, pH 7.4, 150 mM NaCl, 50 mM NaF, 1 mM EDTA, 1% Triton, 5% glycerol) plus 1 μg/ml Leupeptin, 1 mM PMSF, and 1 mM DTT. GST proteins were captured on Pierce glutathione magnetic beads (Thermo Fisher Scientific) for 1 hour, then washed twice in Tris Lysis Buffer, twice in Tris Buffered Saline with 287 mM NaCl and 0.1% Tween 20 (TBST), and twice in TBST with 137 mM NaCl. Each experiment was repeated 3 times, except for K-Ras4A-BRI3BP interaction, which was n=2.

### Western blotting

Primary antibodies: HA tag (CST #3724), GST tag (CST#2625), pERK XP (CST #4370), ERK (CST #4696), GAPDH (CST #2118), MYC tag (CST #2278) Secondary antibodies: Goat Anti Rabbit IgG (H+L) HRP Conjugate (BioRad #172-1019); Goat Anti Mouse IgG (H+L) HRP Conjugate (BioRad #170-6516) Blots (Immobilon-P transfer membrane) were imaged on a ChemiDoc Imager (BioRad) with Clarity or Clarity Max ECL substrates (BioRad 1705060, 1705062). Quantitation was performed in Image Lab Version 6.0 software (BioRad).

### Focus Formation Assay

NIH3T3 cells were transfected with UI4-GFP RNAi expression vectors and US2 vector control or activated Ras expression vector US2-KRasV12. Cells were seeded in 6-well dishes for transfection with a total of 2.5 μg DNA per transfection: 2.25 μg RNAi vector, 0.25 μg US2 or US2-KRasV12. The day following transfection, cells were split to 10 cm dishes (each transfection was split to 2 10-cm dishes) and maintained in DMEM plus 5% calf serum and penicillin-streptomycin for 21 days, with media changed every 4-5 days. Cells were fixed in 10% methanol/10% acetic acid and stained with crystal violet (1% stock solution in 95% ethanol). Foci were quantitated from images captured on a BioRad ChemiDoc imager using the count tool in photoshop (n=2). Similar results were obtained from 3 independent experiments.

pUI4-GFP Luc is a control shRNA expression vector that co-expresses GFP and an shRNA that targets luciferase (50). pUI4-GFP BRI3BP m785 co-expresses GFP and an shRNA that targets mouse BRI3BP. Each vector contains four tandem copies of the shRNA in the miR-155-based SIBR cassette inserted into the second intron of pUI4-GFP (50). The sequences of the predicted mature siRNAs are as follows: Luciferase 1601, 5’-UUUAUGAGGAUCUCUCUGAUUU-3’; BRI3BP RNAi m785, 5’-UGUUAAGCAACCUCACCUGGUU-3’.

### Imaging

For imaging of K-Ras localization, wild type or BRI3BP knockout HeLa cells were transfected with pUS2-GFP-K-RasV12, which expresses green fluorescent protein fused to the N-terminus of activated K-Ras4B (250 ng per well of 12 well) with HeLa Monster (Mirus), per manufacturer’s instructions. pUS2 has been described previously (50). Cells were split to polylysine coated glass coverslips 24 hours after transfection. 48 or 72 hours after transfection, cells were fixed with 4% methanol-free formaldehyde (Thermo #28908) in PBS, permeabilized with 0.1% Triton X-100, stained for GFP (primary antibody, GFP12A6 from Developmental Studies Hybridoma Bank (DSHB); secondary antibody, goat anti-mouse IgG Alexa 488, Thermo #A10680) and nuclei stained with DAPI. DSHB-GFP-12A6 was deposited to the DSHB by DSHB (DSHB Hybridoma Product DSHB-GFP-12A6. For co-localization studies, pUS2-hBRI3BP-mScarlet, which expresses scarlet fluorescent protein (36) fused to the C-terminus of hBRI3BP, was cotransfected with pUS2-GFP-KRasV12 at a 1:1 ratio (500ng) or mCitrine-Rab11a (provided by Joel Swanson, constructed by Xiao-Wei Chen and Alan Saltiel, University of Michigan) at a 1:2 ratio (125ng:250ng) using HeLa Monster. Cells were split to polylysine coated glass coverslips 24 hours after transfection and fixed in 4% formaldehyde at 48 hours. After mounting (Aqua-Poly/Mount, Polysciences, Inc.), cells were imaged on an Olympus FV1000 confocal microscope. Representative images were assembled in Fiji and labeled in Adobe Photoshop. Max intensity images from Z-stacks are shown for GFP-KRasV12 localization, while individual slices are shown for hBRI3BP-mScarlet colocalization. Colocalization analysis was performed on individual slices using EzColocalization in FIJI. GFP-K-RasV12 or mCitrine-Rab11a cells were traced with the elliptical selection tool in FIJI to create regions of interest (ROIs) for analysis of individual cells and Pearson Correlation Coefficients were determined.

## Acknowledgements

We are grateful to Amanda Wilbur for technical assistance, Xu (Jackson) Han for helpful comments in the early stages of this work, and Joel Swanson for discussion of microscopy data. This work was supported by a pilot grant from the University of Michigan Biological Chemistry Department, by the University of Michigan Office of Research, and by the University of Michigan Biomedical Research Council (ABV).

