## Supplemental Figures S1- S6 for "Brain I3 Binding Protein regulates K-Ras4B membrane localization and signaling"

**Supplemental Figure S1**

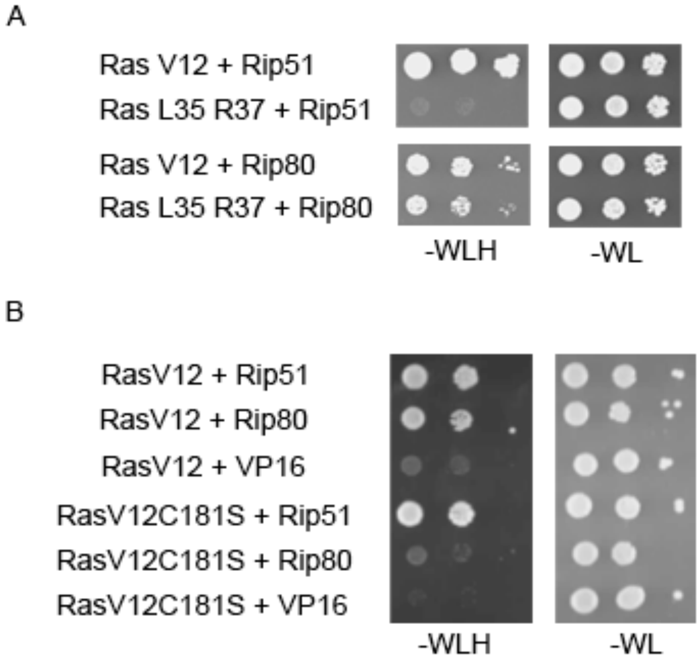


**Figure S1. BRI3BP interacts with Ras in yeast two-hybrid assays.** The L40 yeast reporter strain was transformed with the indicated expression plasmids. Activation of the histidine reporter allows for growth on –WLH plates (minus histidine), indicating a protein-protein interaction. Growth on –WL plates (plus histidine): control. (A) c-Raf/Rip51 (Ras interacting protein 51, two-hybrid isolate) interacts with activated H-Ras V12 but not the effector domain mutant H-Ras L35 R37. In contrast, BRI3BP/Rip80 (Ras interacting protein 80, two-hybrid isolate) interacts with both H-Ras V12 and the H-Ras effector domain mutant. (B) c-Raf/Rip51 interacts with H-Ras, irrespective of prenylation status. BRI3BP/Rip80, in contrast, fails to interact with the prenylation deficient Ras mutant, H-Ras V12 C181S.

**Supplemental Figure S2**

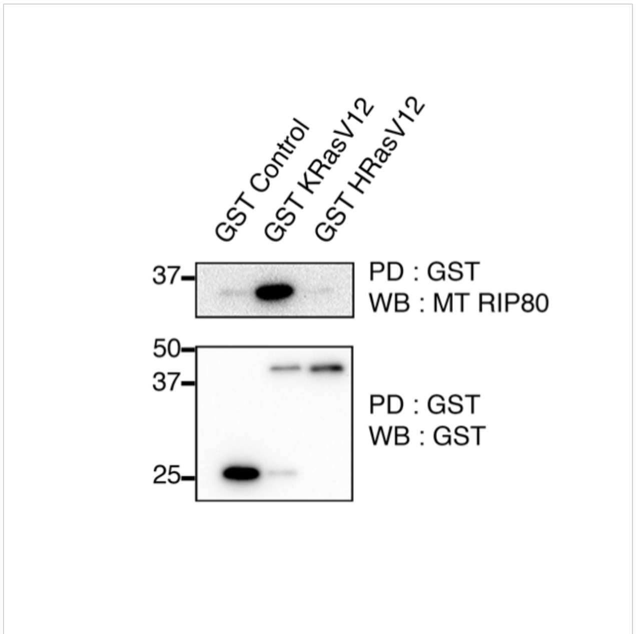


**Figure S2. The BRI3BP Ras interacting domain preferentially interacts with K-Ras in mammalian cells.** GST control or GST-fusion proteins to K-RasV12 or H-RasV12 were co-expressed in HEK cells with the Myc epitope-tagged Ras interacting domain of BRI3BP (MT-Rip80, identified in the yeast two-hybrid screening), by transient expression. GST or GST fusion proteins were isolated from extracts with glutathione agarose beads and co-associated MT-Rip80 was detected by western blot with an antibody to the Myc-epitope tag. BRI3BP/MT-Rip80 preferentially interacts with K-RasV12.

**Supplemental Figure S3**

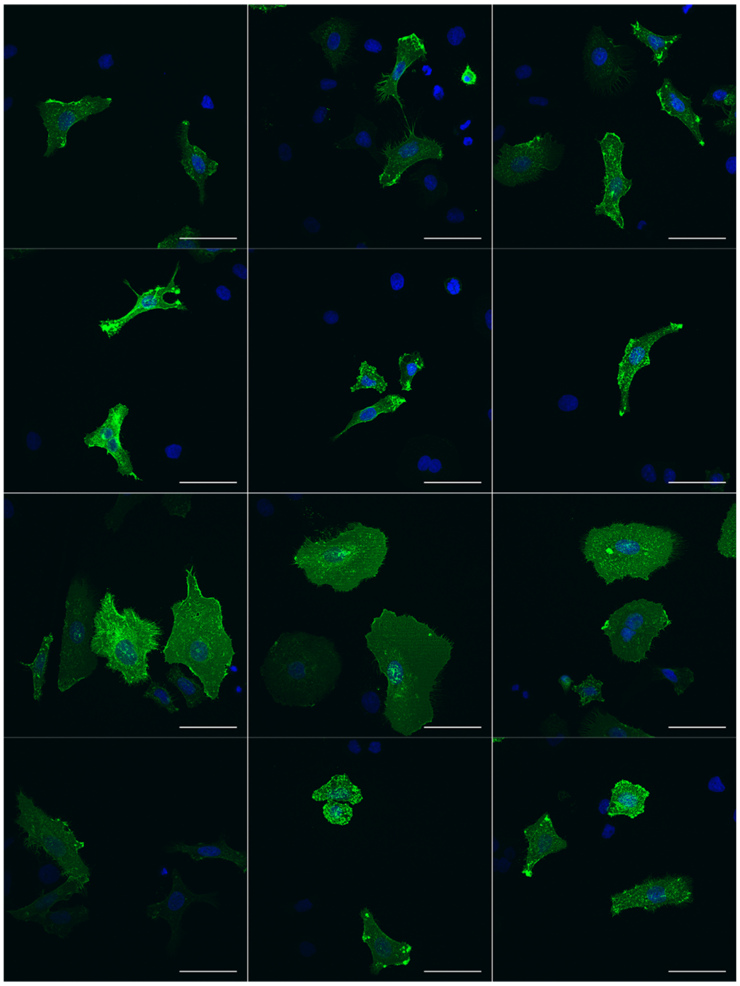


**Figure S3. BRI3BP regulates K-Ras4B plasma membrane localization.** Additional images related to Figure 3. Confocal microscopy of GFP-K-RasV12 localization in wild type HeLa cells (WT, row 1), wild type cells with overexpressed BRI3BP (WT + BRI3BP, row 2), CRISPR/Cas9 BRI3BP knockout cells (KO, row3), or knockout cells with ectopic BRI3BP (complementation) (KO + BRI3BP, row 4). Scale bars: 50 μm. Cells were fixed 48 hours after transfection and processed for indirect immunofluorescent detection of GFP.

**Supplemental Figure S4**

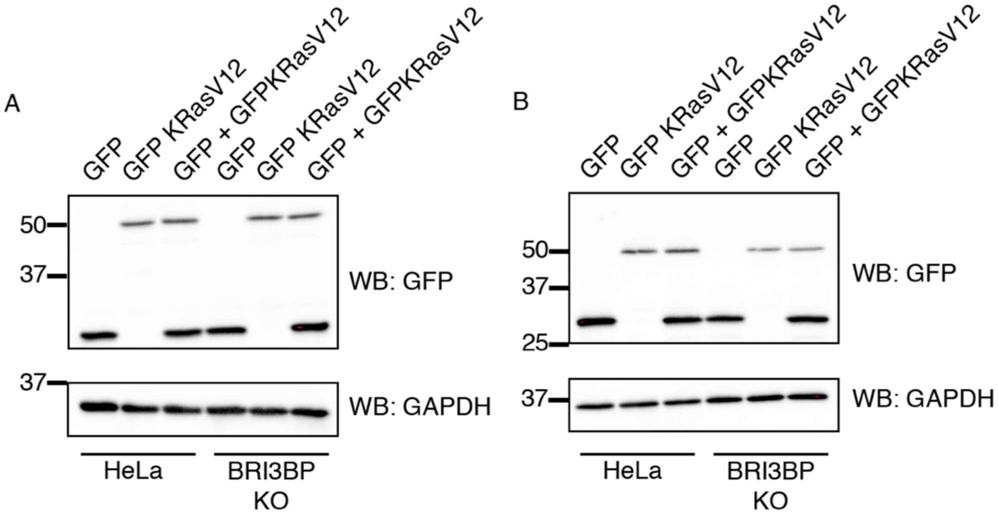


**Figure S4. Western blot showing levels of GFP-K-RasV12 in wild type or BRI3BP knockout HeLa cells.** GFP, GFP-K-RasV12, GFP-K-RasV12 plus GFP expression plasmids were transfected into HeLa WT or BRI3BP knockout cells. Extracts were prepared 48 hours (A) or 72 hours (B) after transfection and GFP or GFP-K-RasV12 protein detected by western blot with an antibody to GFP after SDS-PAGE. Wild type and BRI3BP knockout cells have similar levels of GFP-K-RasV12 protein at both 48 and 72 hours.

**Supplemental Figure S5**

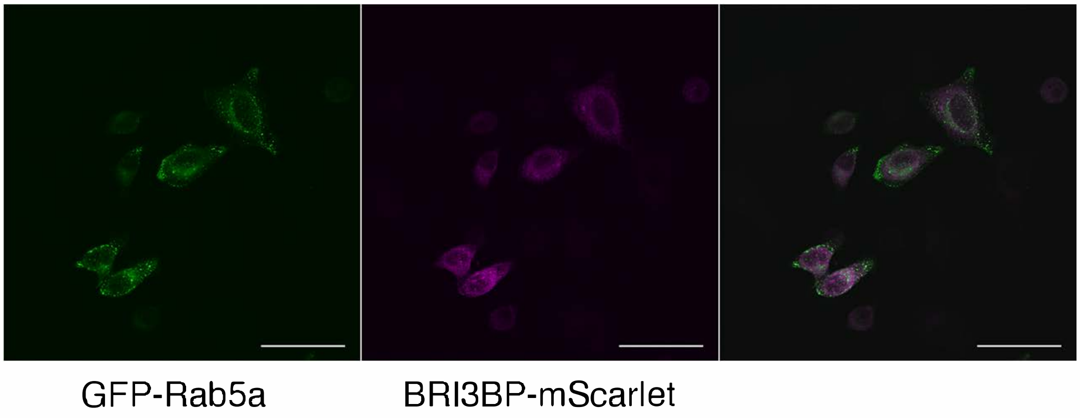


**Figure S5. BRI3BP does not co-localize with the early endosomal marker Rab5a.** BRI3BP-mScarlet and GFP-Rab5a were co-transfected (2:1 ratio) into HeLa cells. Cells were fixed for imaging 48 hours after transfection.

**Supplemental Figure S6**

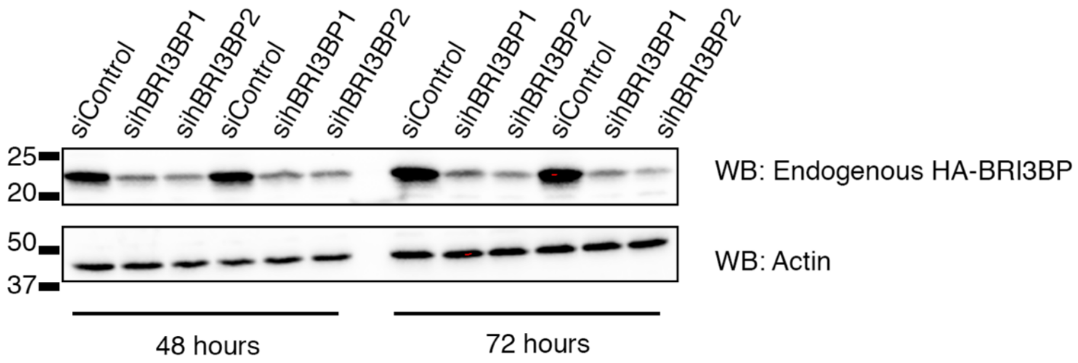


**Figure S6. BRI3BP-HA knockdown by siRNAs.** Western blot for endogenous BRI3BP-HA (upper panel). Actin (loading control): lower panel. HEK cells with endogenous C-terminal HA epitope tagged BRI3BP were transfected with a control or one of two BRI3BP siRNAs, targeting different sequences (10 nM). Anti-HA western blot analysis of extracts prepared 48 hours after transfection shows decreased HA-BRI3BP in BRI3BP siRNA transfected samples relative to control.
